# Development, feasibility and potential effectiveness of community-based continuous mass dog vaccination delivery strategies: lessons for optimization and replication

**DOI:** 10.1101/2022.03.10.483887

**Authors:** Christian Tetteh Duamor, Katie Hampson, Felix Lankester, Ahmed Lugelo, Emmanuel Mpolya, Katharina Kreppel, Sarah Cleaveland, Sally Wyke

**Affiliations:** Department of Global Health, Nelson Mandela African Institute of Science and Technology, Arusha – Tanzania; Environmental Health and Ecological Sciences Thematic Group, Ifakara Health Institute – Tanzania; Institute of Biodiversity, Animal Health & Comparative Medicine, College of Medical, Veterinary and Life Sciences, University of Glasgow, Glasgow, G12 8QQ – UK; Paul G. Allen School for Global Animal Health, Washington State University, 240 SE Ott Road, Pullman, WA 99164-7090 – USA; Serengeti Health Initiative, Serengeti – Tanzania; Sokoine University of Agriculture, Morongoro – Tanzania; Institute of Health and Wellbeing, College of Social Sciences, University of Glasgow, Glasgow, G12 8QQ – UK

**Keywords:** community-based, dog vaccination, process evaluation, rabies

## Abstract

**Objectives:** Dog vaccination can eliminate rabies, but annual delivery strategies do not sustain vaccination coverage between campaigns. We describe the development of a community-based continuous mass dog vaccination (CBC-MDV) approach designed to improve and maintain vaccination coverage in Tanzania and examine the feasibility of delivering this approach as well as lessons for its optimization.

**Methods:** We developed three delivery strategies of CBC-MDV and tested them against the current annual vaccination strategy following the UK MRC’s guidance: i) developing an evidence-based theoretical framework of intervention pathways and ii) piloting to test feasibility and inform optimization. For our process evaluation of CBC-MDV we collected data using non-participant observations, meeting reports and implementation audits and in-depth interviews, as well as household surveys of vaccination coverage to assess potential effectiveness. We analyzed qualitative data thematically and quantitative data descriptively.

**Results:** The final design included delivery by veterinary teams supported by village-level one health champions. In terms of feasibility, we found that less than half of CBC-MDV’s components were implemented as planned. Fidelity of delivery was influenced by the strategy design, implementer availability and appreciation of value intervention components, and local environmental and socioeconomic events (e.g. elections, funerals, school cycles). CBC-MDV activities decreased sharply after initial campaigns, partly due to lack of supervision. Community engagement and involvement was not strong. Nonetheless, the CBC-MDV approaches achieved vaccination coverage above the critical threshold (40%) all-year-round. CBC-MDV components such as identifying vaccinated dogs, which village members work as one health champions and how provision of continuous vaccination is implemented need further optimization prior to scale up.

**Interpretation:** CBC-MDV is feasible to deliver and can achieve good vaccination coverage. Community involvement in the development of CBC-MDV, to better tailor components to contextual situations, and improved supervision of activities are likely to improve vaccination coverage in future.

**AUTHOR SUMMARY:** Annual mass dog vaccination campaigns that reach at least 70% of the dog population, should maintain sufficient herd immunity between campaigns to interrupt rabies transmission. However, it is often challenging to reach 70% of the dog population with annual vaccination campaigns. We hypothesized that a community-based continuous approach to dog vaccination could better maintain high levels of vaccination coverage all-year-round. We describe the development of a community-based continuous approach to dog vaccination in Tanzania, and assessed the feasibility of delivering its components, its potential effectiveness and lessons for its optimization. We found that the approach was well accepted, as its development involved key stakeholders. Although less than half of the components of the community-based continuous approach were delivered exactly as planned, over 70% of dogs were vaccinated and the approach maintained coverage above the critical vaccination threshold throughout the year. We conclude that it is feasible to deliver a community-based continuous approach to dog vaccination, but that some components need further improvement; more supervision and community involvement should lead to better outcomes.

## INTRODUCTION

Rabies is a central nervous system infection that can infect all mammals. The disease has a case fatality rate approaching 100% [1,2]. Globally, human deaths are estimated at about 59,000 per annum, with 99% due to domestic dog-mediated transmission [3]. The burden of rabies is highest in endemic regions where both human and animal rabies vaccines are not reliably accessible [3].

Rabies is controllable for several reasons: domestic dogs are the primary source of infections to humans and rabies has a consistently low basic reproductive number (R_0_<2) across a wide range of settings [1]; dogs in endemic regions are typically accessible for vaccination [4,5]; and the low R_0_ means that the critical vaccination threshold required to achieve herd immunity is relatively low (approximately 40%). Despite these reasons, rabies remains endemic in many settings and only limited dog vaccination is undertaken. A possible concern should mass dog vaccination be scaled up is that, despite the critical vaccination threshold being low, to sustain vaccination coverage *above* this level over the course of the year, annual vaccination campaigns must reach a higher proportion of dogs, of around 70% [1,6,7].

Most endemic countries where some mass dog vaccination (MDV) has been initiated, including Tanzania, use annual team-delivered approaches in which government vaccination teams use cold-chain stored vaccines to conduct annual vaccination clinics in targeted villages. However, annual team-delivered campaigns (subsequently referred to in this study as the pulse approach) are affected by several factors that limit their ability to achieve and sustain vaccination coverages above the critical threshold to control rabies. These include: high rates of dog population turnover in most endemic countries, which results in rapid declines in population immunity in the interval between annual campaigns [8,9]; teams needing to travel long distances on dirt roads from cold chain facilities, which is sometimes possible only at certain times of the year; campaign day(s) being negatively affected by agricultural cycles, inclement weather, school days, funerals, and local festivals [7]; high fixed vehicle and personnel costs, with the cost-per-dog vaccinated reaching as high as $7.36 [10–12].

Recent research has shown that Nobivac Canine Rabies Vaccine, a widely used vaccine for dogs [13], is thermostable and can induce equivalent immune responses following non cold chain storage at temperatures up to 30°C for three months. Thermotolerance remained when the vaccines were stored in rural Tanzania in locally made passive cooling devices, within which temperatures were kept relatively cool despite ambient temperatures reaching 37°C [14]. This research has created opportunities for new approaches to rabies vaccine distribution and delivery, including options for the storage of vaccines in remote communities which would allow all year-round routine vaccination of dogs by community-based personnel. A community-based continuous mass dog vaccination (CBC-MDV) approach has the potential to sustain population immunity above the critical threshold, as new puppies and other susceptible dogs (for example, newly acquired dogs or those that missed previous vaccination campaigns) can be vaccinated without having to wait for the annual campaign. Through empowering communities to own and sustain local dog vaccination efforts, it has been hypothesized that a CBC-MDV model could also result in more dogs being reached at less cost per animal vaccinated [13,15,16].

CBC-MDV is a complex intervention, with several interacting components such as the involvement of local veterinary authorities and communities, local storage of dog rabies vaccines outside of the cold chain system and a continuous approach to dog vaccine delivery. Consequently, the intervention could operate differently in different settings. The UK Medical Research Council Guidance on developing and evaluating complex interventions prior to full scale evaluation recommends a systematic approach to intervention development [17]. This approach should include the development of the intervention with stakeholders, a theoretical understanding of how it is likely to operate, and piloting of its delivery with a view to evaluating the feasibility of its delivery in the long term, prior to full scale evaluation. The study describes formative work that took place to design a model of CBC-MDV and the development and testing of three different delivery strategies for CBC-MDV. Following the UK Medical Research Council Guidance, we describe the development and evaluation of the feasibility of delivering CBC-MDV. We also assess the potential of CBC-MDV to sustain vaccination levels above the critical threshold for rabies elimination and lessons for its optimization and replication.

## METHODS

The research was conducted in two stages, Figure 1 provides a schematic overview of the processes involved.

**Fig 1.**
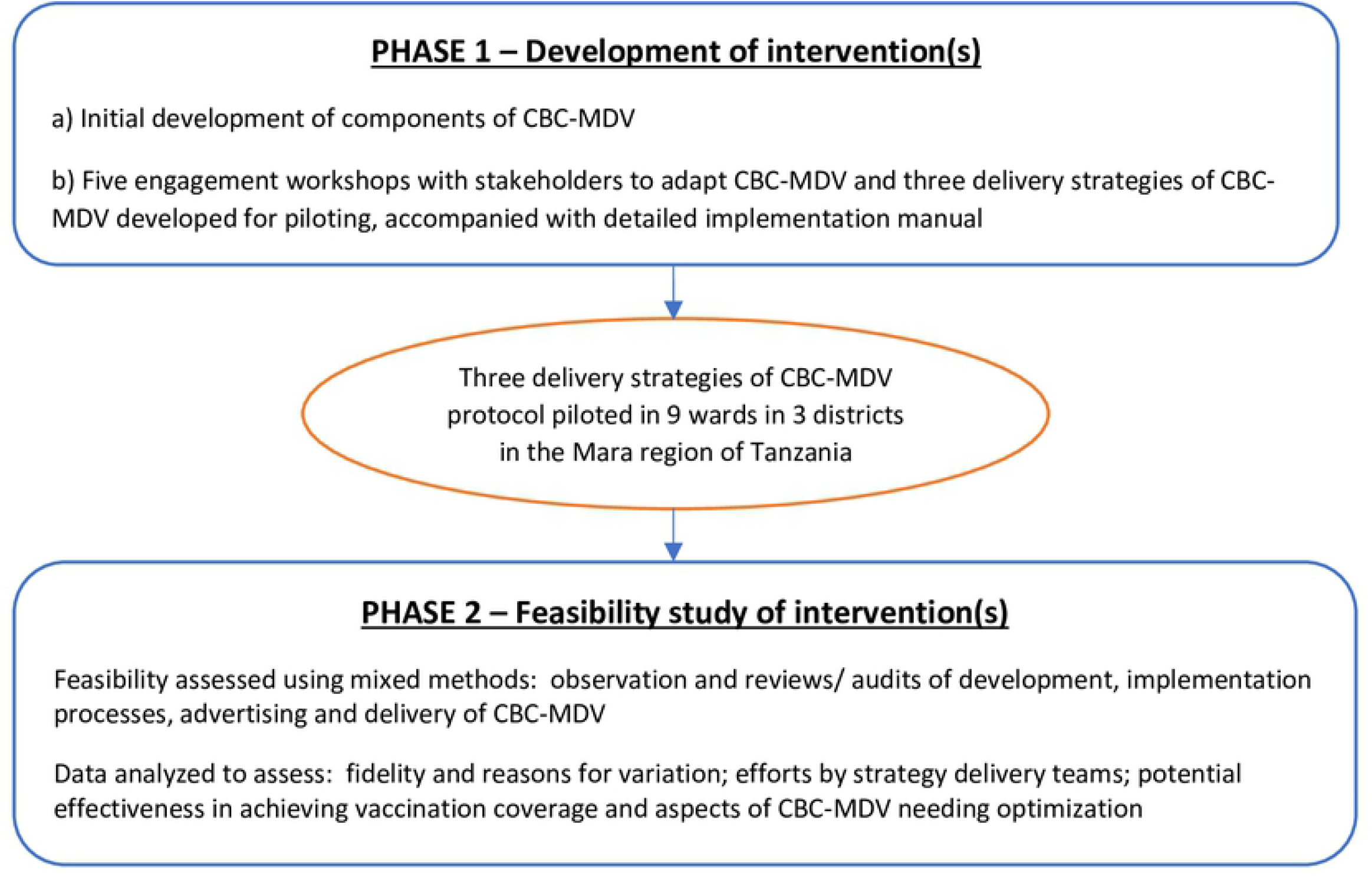
Development and optimization process for Community-based Continuous Mass Dog Vaccination (CBC-MDV) prior to full-scale evaluation

### Phase 1: Developing components of CBC-MDV

CBC-MDV was developed to be delivered in rural Tanzania and piloted in three districts of Rorya, Tarime and Butiama of the Mara region, north-west Tanzania between Lake Victoria and Kenya. This area is home to several ethnic groups who are primarily engaged in agro-pastoral and fishing activities. Dog ownership is common with larger households and those having livestock tending to own more dogs [18–20]. The pilot phase included 12 wards drawn from each district: wards are a cluster of 3-4 villages; villages are divided into subvillages; the number of subvillages per village ranged from 2 to 13 in our study area.

Evidence on barriers to a centralized, team-delivered dog vaccination approach (as laid out in the introduction), the feasibility of storing the Nobivac Rabies Vaccine in locally made passive cooling devices [13,14] and the ability of community-based persons to vaccinate dogs [21] provided the context for developing initial components of CBC-MDV.

The initial design was discussed with potential stakeholders in the Mara region (where a large-scale randomized controlled trial (RCT) is proposed to take place following on from this pilot study) and subsequently with national level veterinary officials and international experts, with workshops taking place between May 2018 and May 2019. Table 1 describes the stakeholder groups involved and aim of each workshop.

**Table 1.** Stakeholder group, purpose and date of engagement workshops

|  | **Stakeholder Group** | **Purpose of Workshop** | **Dates; Venue** |
| --- | --- | --- | --- |
| 1 | National Level Veterinary Officials, Mara Regional Medical and Veterinary Officers, District Medical and Veterinary Officers, Nurses and Livestock Field Officers plus research staff | To introduce national veterinary officials and Mara region stakeholders to potential CBC-MDV strategies | 23-26 May, 2018;<br>Mugumu – Serengeti |
| 2 | Veterinary technical staff from Ministry of Livestock and Fisheries Development, community health specialist from World Health Organization – Tanzania country office, the Mara Regional Medical Officer, representatives from Ministry of Health and Tanzanian One Health Coordination Unit plus research staff. | To share evidence for the safety of use of locally made passive cooling devices to store vaccines & non-animal health professionals to vaccinate dogs and to demonstrate that the research evidence was strong enough for local use. | 17- 18 July, 2018;<br>The Prime Minister’s Office – Dar es Salaam |
| 3 | Three Rabies Researchers from Global Animal Health – Tanzania, Director of Veterinary Services and Registrar of Tanzanian Veterinary Council | To provide the outcome of Workshop 2, and to share evidence of use of locally made passive cooling devices to store vaccines & non-animal health professionals to vaccinate dogs | 17th November, 2018;<br>Office of Director of Veterinary Services – Dodoma |
| 4 | Researchers from Washington State University (5), University of Glasgow (5), Global Animal Health – Tanzania (6), Director of Veterinary Services, Chairman and Registrar of Tanzania Veterinary Council, President of Tanzania Veterinary Association, representatives from Ministry of Health and One Health Coordination Unit | To finalize design of CBC-MDV for the pilot study, define roles of district, ward and village level implementers and to launch the research project | 22nd – 23rd Mar, 2019;<br>Arusha. |
| 5 | Mara Regional Commissioner and Administrative Secretary, Researchers from Global Animal Health – Tanzania (6), Director of Veterinary Services, Chairman and Registrar of Tanzania Veterinary Council, President of Tanzania Veterinary Association, Mara Regional Medical and Veterinary Officers, District Medical and Veterinary Officers, Nurses and Livestock Field Officers | To bring the research team and human and animal health staff of the Mara region together, to outline logistical needs for implementing CBC-MDV and to declare the research as a national learning project to inform national mass dog vaccination strategies for Tanzania | 7th – 8th May, 2019;<br>Office of the Mara Regional Commissioner |

The first author participated in and made notes (11 observation days) of all the workshops, and documented stakeholders’ opinions and concerns of CBC-MDV, specifically: how vaccines will be stored outside of the cold chain system in wards using locally made passive cooling devices, the level of training required to vaccinate dogs, local involvement in implementation and roles at district, ward and village levels. The research team met after each workshop to revise the components of CBC-MDV.

Following the final workshop, the research team developed a theory of change model and a manual to guide implementers (district livestock field officers, ward-based rabies coordinators – RCs and village-based one-health champions – OHCs) in delivering the CBC-MDV components. To identify the most efficient approach to delivering the components, three delivery strategies of CBC-MDV were designed to be piloted.

### Phase 2: Feasibility of delivering CBC-MDV, potential effectiveness and lessons learned

The three delivery strategies of CBC-MDV were piloted over a 12-month period and evaluated using mixed methods and the outcomes compared to that of the pulse (annual team-delivered) approach which was also undertaken as part of the pilot study. Table 2 summarizes which methods were used to assess the feasibility and potential effectiveness of the delivery strategies as well as to formulate lessons learned.

**Table 2.**
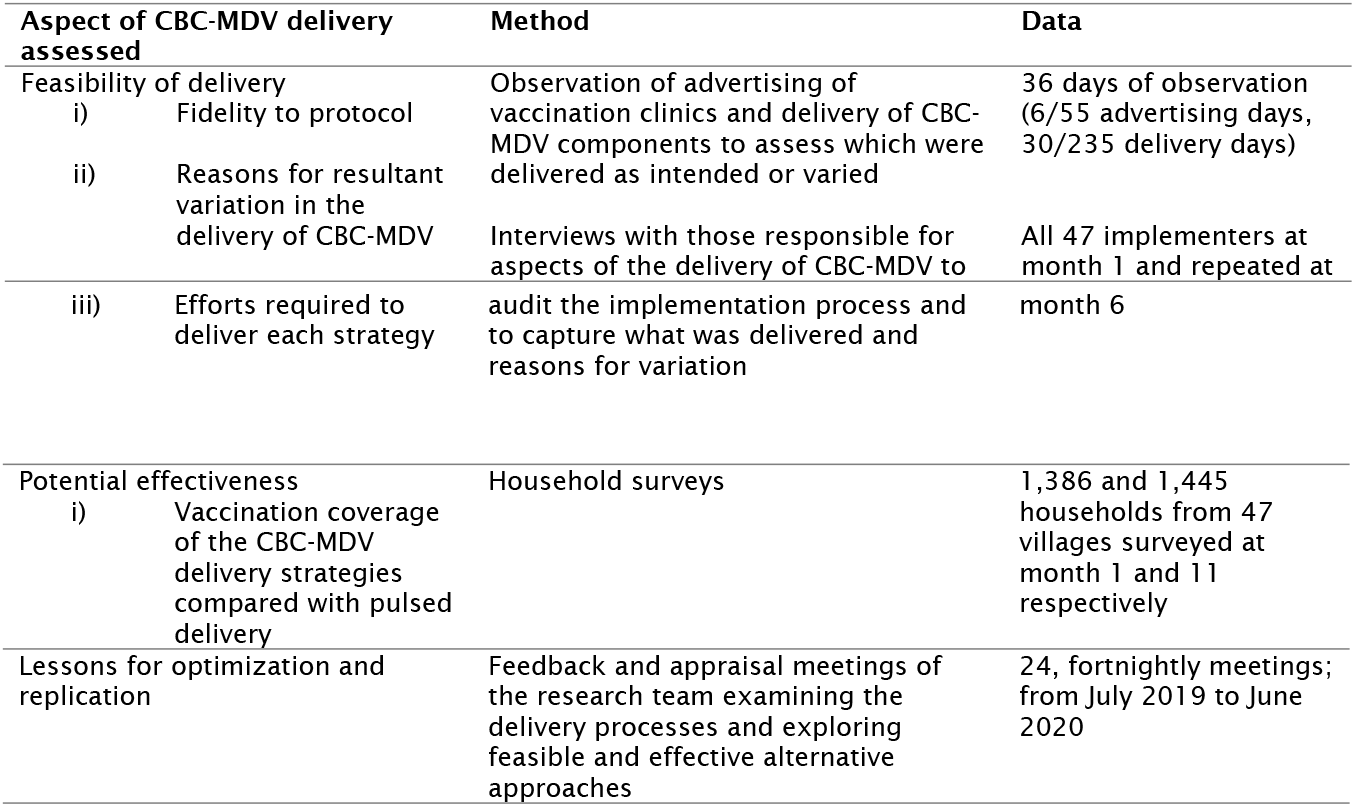
Summary of research methods used to assess the feasibility of delivering community based continuous mass dog vaccination (CBC-MDV), potential effectiveness and formulate lessons learnt

#### Assessing Fidelity, Variation and Efforts

To assess the fidelity of the implementation process during phase 2 and the reasons for variation in delivering CBC-MDV, we conducted observations on advertising campaigns (6/55 days) and delivery of vaccination activities (30/235 days) noting whether implementers delivered components of CBC-MDV as planned and factors responsible for variation.

We audited delivery of CBC-MDV using semi-structured interviews with implementers (one with each of the 47 implementers) about aspects of delivery, record review, inspection of how vaccines were managed at district veterinary offices and wards, and installation and maintenance of locally made passive cooling devices and their temperature loggers within wards. Notes were taken on which components of CBC-MDV were delivered as planned and on potential reasons for variation. The audits were carried out early in the delivery of CBC-MDV at month 1 and repeated at month 6.

We used observation and audit data to assess and compare efforts required for each of the CBC-MDV strategies and the fidelity of their delivery.

#### Assessing potential effectiveness

When dogs were vaccinated owners were given a vaccination certificate and dogs were microchipped. To assess how the strategies performed with respect to vaccination coverage, random samples of households (Table 2) were surveyed in each village, scanning dogs for a microchip and inspecting vaccination certificates. If neither the dog nor the certificate could be found, we asked household members whether their dog(s) had been vaccinated. The surveys were conducted at month 1 and 11 after roll out of CBC-MDV. Detailed reports of outcome measurement are presented in an outcome evaluation paper (Lugelo et al. preprint) and are summarized in this manuscript to provide informative context to the process evaluation.

#### Lessons for optimization and replication

To optimize CBC-MDV, the research team reviewed the observation and audit data on the delivery process through fortnightly feedback and appraisal meetings to identify components of CBC-MDV that were not working and designed alternative approaches. The team also identified best practices by implementers and components of CBC-MDV that were context sensitive. The first author participated in these meetings and made detailed reports.

### Data analysis

#### Fidelity, Variation and Effort

To assess the extent to which the components were delivered as intended, field notes from observations of advertising and from the audits of the implementation process were read and summarized as either ‘delivered as planned’, ‘delivery modified’, ‘not delivered as planned’ or ‘delivered in excess of what was planned’. To assess the reasons for variation from what was planned, qualitative notes from observation of the advertising process and audits were thematically analysed as follows. The first author developed the initial coding frame using a combination of deductive and inductive approaches [22,23]. Two authors independently applied the coding frame to a sample of the data (2 observation and 1 audit notes), and the coding frame was discussed and amended over three iterations. The first author then applied the coding frame to the whole data set. The main themes were: community engagement, estimation of dog population, advertising of campaigns, starting and closing time of vaccination clinics, delivery of continuous vaccination and choice of approaches for clinics. The coded texts were used in complementing, expanding and elaborating on understanding of the manner in which CBC-MDV was delivered and factors that influenced feasibility of delivering the different components. Qualitative data analysis was done using QSR NVIVO version 12.5.0 (NVivo qualitative data analysis software; QSR International Pty Ltd. Version 12, 2018).

To assess the effort that was required to implement each of the three CBC-MDV strategies data were collected on the number of times and hours spent advertising, and number of campaigns delivered. These data were examined to determine whether the efforts varied by strategy after plotting in Excel version 16.

#### Assessing potential effectiveness

Vaccination coverage achieved by each delivery strategy was calculated as the proportion of the dog population surveyed that had either i) a microchip, ii) a vaccination certificate or iii) owner recall that the dog had been vaccinated. We summarized the coverage estimates at month 1 and month 11, annual averages achieved by each CBC-MDV strategy and the pulse strategy.

#### Lessons for optimization and replication

To ensure successful replication of CBC-MDV in other settings, the research team, through the appraisal meetings, identified components of CBC-MDV that were appreciably influenced by contextual factors. Reference was made to the literature on how certain barriers to implementing community-based interventions were overcome and considered in optimizing CBC-MDV. Based on the conclusions reached by the research team, alternative approaches were designed for the CBC-MDV components that were not working as planned. Best practices among implementers were identified and incorporated into the CBC-MDV design for implementation in the full-scale trial planned for the Mara region.

### Ethics statement

The study was approved by the Institutional Animal Care and Use Committee, Washington State University [Approval No. 04577 – 001], the Tanzania National Medical Research Institute [NIMR/HQ/R.8a/Vol.IX/2788] and Ifakara Health Institute [IHI/IRB/No:024-2018].

## RESULTS

### Phase 1: Development of CBC-MDV intervention

Table 3 summarizes the essential components of CBC-MDV, the rationale for their inclusion, the views on each component expressed by stakeholders during meetings and adaptations made to the design of the components to address concerns. The detailed components of each ingredient are outlined in Supplementary Table 1. The development process of CBC-MDV was iterative and participation in the workshops was multisectoral and included participants who both work in either the public health or animal health sector and are members of local communities, but did not specifically include community leaders/ decision-makers.

**Table 3.**
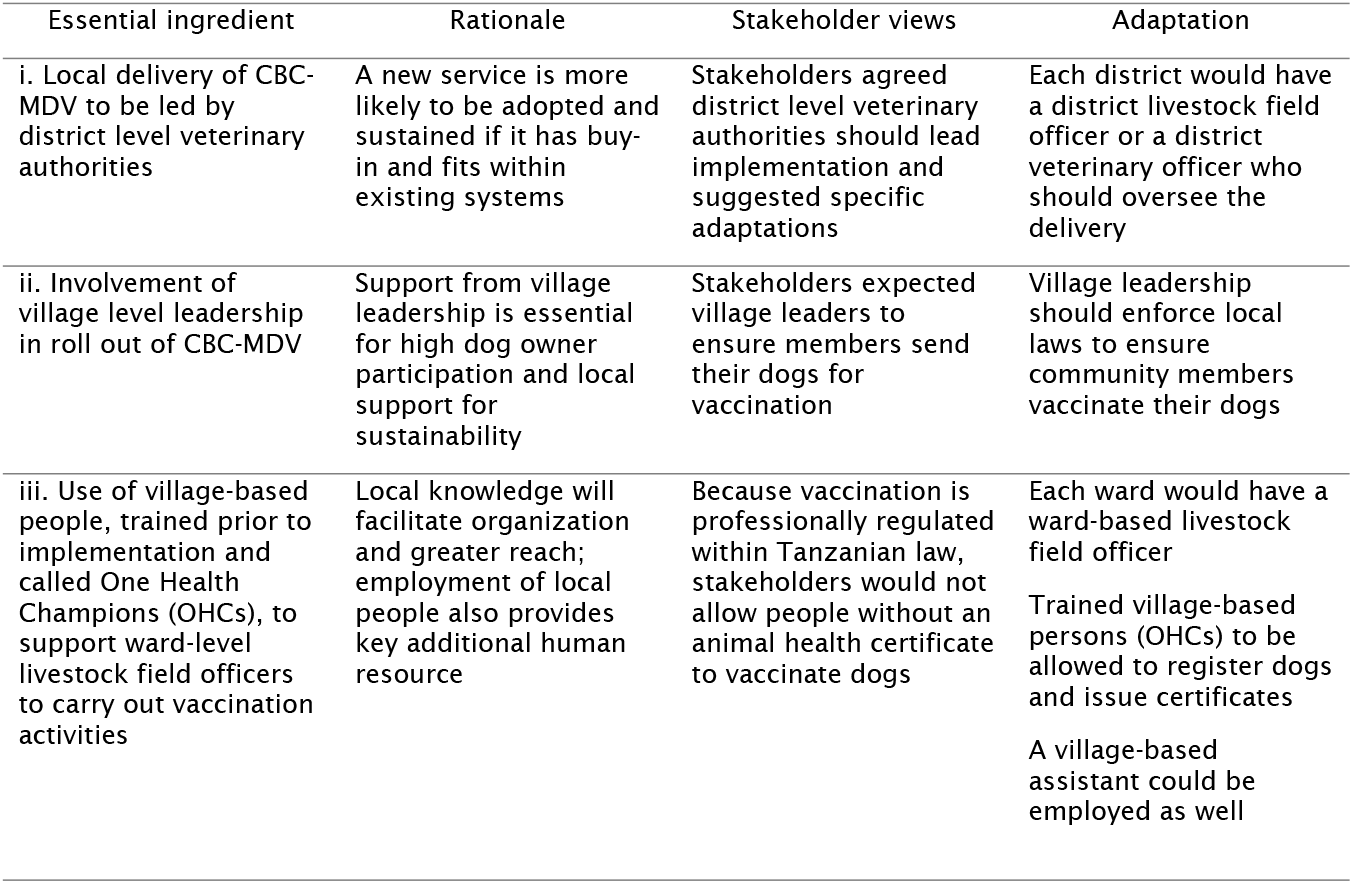
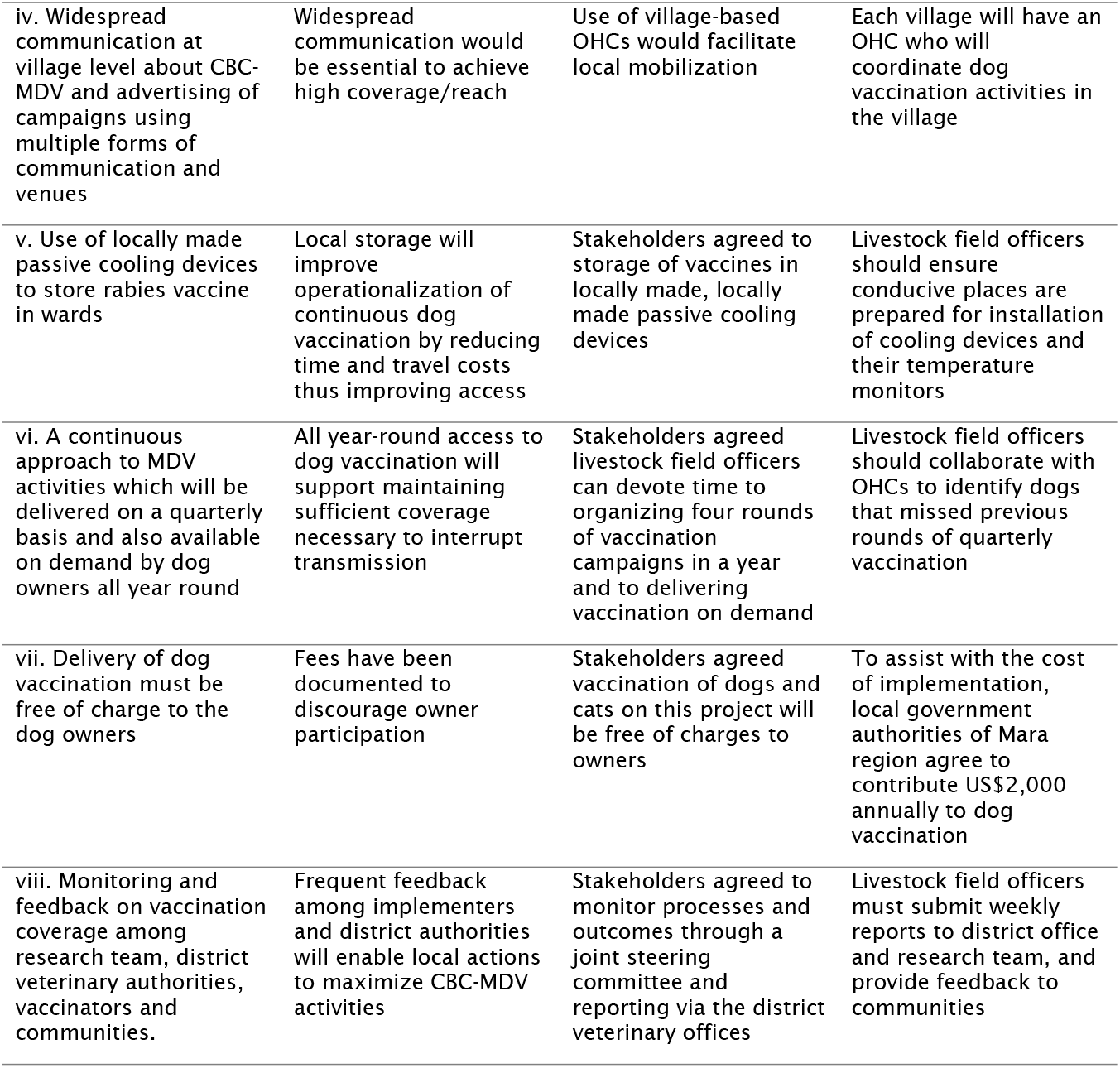
Essential components of CBC-MDV and responses to stakeholder concerns.

| Essential ingredient | Rationale | Stakeholder views | Adaptation |
| --- | --- | --- | --- |
| i. Local delivery of CBC-MDV to be led by district level veterinary authorities | A new service is more likely to be adopted and sustained if it has buy-in and fits within existing systems | Stakeholders agreed district level veterinary authorities should lead implementation and suggested specific adaptations | Each district would have a district livestock field officer or a district veterinary officer who should oversee the delivery |
| ii. Involvement of village level leadership in roll out of CBC-MDV | Support from village leadership is essential for high dog owner participation and local support for sustainability | Stakeholders expected village leaders to ensure members send their dogs for vaccination | Village leadership should enforce local laws to ensure community members vaccinate their dogs |
| iii. Use of village-based people, trained prior to implementation and called One Health Champions (OHCs), to support ward-level livestock field officers to carry out vaccination activities | Local knowledge will facilitate organization and greater reach; employment of local people also provides key additional human resource | Because vaccination is professionally regulated within Tanzanian law, stakeholders would not allow people without an animal health certificate to vaccinate dogs | Each ward would have a ward-based livestock field officer<br>Trained village-based persons (OHCs) to be allowed to register dogs and issue certificates<br>A village-based assistant could be employed as well |
| iv. Widespread communication at village level about CBC-MDV and advertising of campaigns using multiple forms of communication and venues | Widespread communication would be essential to achieve high coverage/reach | Use of village-based OHCs would facilitate local mobilization | Each village will have an OHC who will coordinate dog vaccination activities in the village |
| v. Use of locally made passive cooling devices to store rabies vaccine in wards | Local storage will improve operationalization of continuous dog vaccination by reducing time and travel costs thus improving access | Stakeholders agreed to storage of vaccines in locally made, locally made passive cooling devices | Livestock field officers should ensure conducive places are prepared for installation of cooling devices and their temperature monitors |
| vi. A continuous approach to MDV activities which will be delivered on a quarterly basis and also available on demand by dog owners all year round | All year-round access to dog vaccination will support maintaining sufficient coverage necessary to interrupt transmission | Stakeholders agreed livestock field officers can devote time to organizing four rounds of vaccination campaigns in a year and to delivering vaccination on demand | Livestock field officers should collaborate with OHCs to identify dogs that missed previous rounds of quarterly vaccination |
| vii. Delivery of dog vaccination must be free of charge to the dog owners | Fees have been documented to discourage owner participation | Stakeholders agreed vaccination of dogs and cats on this project will be free of charges to owners | To assist with the cost of implementation, local government authorities of Mara region agree to contribute US\$2,000 annually to dog vaccination |
| viii. Monitoring and feedback on vaccination coverage among research team, district veterinary authorities, vaccinators and communities. | Frequent feedback among implementers and district authorities will enable local actions to maximize CBC-MDV activities | Stakeholders agreed to monitor processes and outcomes through a joint steering committee and reporting via the district veterinary offices | Livestock field officers must submit weekly reports to district office and research team, and provide feedback to communities |

### The strategies of CBC-MDV tested

Stakeholders determined that the essential components of CBC-MDV could be delivered slightly differently and used the pilot (phase 2) to assess the three forms of delivery (Table 4), each of which included the essential components. A ward from each district was allocated to each of the three CBC-MDV delivery strategies. An additional ward from each district was then allocated to the pulse (once annual) strategy.

**Table 4.** Strategies for delivering components of CBC-MDV in the pilot study

| Strategy | Frequency | Rationale |
| --- | --- | --- |
| **One: Village level** temporal static point clinics in month 1 for all villages in the ward | Campaigns repeated at months 3, 6, and 9 using either the same approach or house-to-house, plus on-demand vaccination, i.e. responding to alerts from owners of dogs needing vaccination | Within three months enough puppies and new dogs would have arrived in villages in manageable numbers for efficient vaccination |
| **Two: Subvillage level** temporal static point clinics in month 1 for all villages in the ward. | Campaigns repeated at months 3, 6, and 9 using same approach or house-to-house, plus on-demand. | Bringing clinic centers closer to more people should increase owner participation |
| **Three:** Implementers will deliver mass dog vaccinations using whichever of the above methods they consider to be best. | Continuous quarterly campaigns (at months 1, 3, 6, and 9) | Discretion to implementers and their knowledge of local terrain and context will influence their choices of vaccination approach and improve performance. |

Figure 2 presents the logic model agreed between research team members and the stakeholder groups as to how CBC-MDV in general is expected to work.

### Phase 2: Assessment of feasibility and potential effectiveness

#### Fidelity and Reasons for Variation

Table 1 (supplementary file) presents an expanded form of the essential (45) components of CBC-MDV and summary analysis of fidelity of delivery: 20 components (44%) were delivered as planned, 14 (31%) were not delivered at all, nine (20%) were modified and two (5%) were delivered in excess of what was planned. The components were broadly categorized into eight groups (as detailed in Table 3) and their fidelity described as follows:

##### i. Local delivery of CBC-MDV to be led by district level veterinary authorities to foster buy-in

Of the four components relating to district veterinary authority roles, two were modified in delivery. To foster community acceptance of the one health champions and rabies coordinators, the district livestock field officers were to write letters to introduce the vaccinators to their villages. All the district officers wrote letters after the training workshop. The district officers took stocks of vaccines received from the research project and distributed them to wards as planned. However, vaccines returned from two wards to district offices were not labeled and stored as planned. The district officials reported only supervising and monitoring campaigns as part of routine district veterinary functions. They cited lack of vehicle and fuel as key challenges to supervision. All the RCs reported they were not supervised by district officials as planned.

##### ii. Involvement of village level leadership in roll out of CBC-MDV to foster owner participation and local support

There were five components of CBC-MDV to be implemented to bring community leadership on-board with delivery of dog vaccination. Of these, four were modified or partly delivered as planned and one was not delivered. Of 35 OHCs, the majority received letters introducing them to their villages (31, 89%). However, most of them received the letters just a few days before or after the process had started and there were very few or no opportunities to introduce them at village meetings. Of the 19 (54%) introduced, 17 were introduced only in a leaders’ meeting; while in the cases of those not introduced (16, 46%), the RCs or OHCs only informed ward or village executive officers about the programme. Hence, most villagers did not have the opportunity to link the RCs and OHCs with the vaccination campaigns before they started.

The protocol also required RCs to discuss vaccination timetables with village leaders; only four out of nine RCs reported directly informing a community leader about their timetables. Again, OHCs were to work with ‘mabalozi’ (leaders of a cluster of ten houses) to estimate the village dog population. These were partly implemented; only a few (3, 9%) OHCs reported working with ‘mabalozi’; the rest either went to houses directly (19, 54%) or instead worked with subvillage chairpersons (13, 37%). The frequently cited reasons for not working with ‘mabalozi’ included: ‘mabalozi’ perceived OHCs as not belonging to their political party or seen the project as not a community agenda and hence requested money (15, 43%); *“one ‘balozi’ said, you went to the workshop and received big allowances and you have come to tell us to go and work”* [OHC, Implementation Audit, Strategy 1-Tarime]. Also, the concept of ‘mabalozi’ is not practiced uniformly across all jurisdictions (11, 31%). Other reasons were OHCs thought they were to work instead with subvillage chairpersons (6, 17%) or they did not trust ‘mabalozi’ to produce accurate figures on the dog population (4, 11%).

##### iii. Use of trained village-based One Health Champions to support ward-level rabies coordinators with local knowledge to carry out vaccination activities

There were six essential ingredients relating to village-based personnel supporting delivery of CBC-MDV at village levels. Out of these six, two were delivered as planned, one was partly delivered, two were not implemented and one was implemented in excess of what was planned. To ensure that only the required number of vaccines for a round were requested, all OHCs (35, 100%) provided estimates of the village dog population to RCs for request of vaccination materials. All OHCs also advertised vaccination clinics as planned. On the other hand, only two out of 35 OHCs conducted sensitization in village meetings. The opportunities for OHCs were likely limited as most of the villages did not hold meetings before the start of campaigns. Over the course of the year none of the OHCs documented dogs that missed the previous rounds as planned. All OHCs supported vaccination clinics in other villages of the ward in addition to theirs, as the workload at a center is ideally for three people. Not all of OHCs had cooperation from their village leadership, possibly because most of the OHCs were not persons with influential village positions.

##### iv. Widespread communication at village level about CBC-MDV and advertising of campaigns using multiple forms of communication and venues to promote high reach

Advertising of campaigns was largely carried out as planned. Of three components relating to advertising, one was delivered as planned, one modified and one delivered in excess of what was planned. All OHCs (35, 100%) delivered the complete contents of the adverts as designed, which included: date, time, location of clinic, specified animals to be vaccinated as dogs and cats, and vaccination being free-of-charge, using mega phones and posters at vantage points as prescribed. However, instead of the night before, announcements started two to three days before, likely occasioned by perceived workload (nature of settlement and size of villages – need to cover long distances). Out of a total of 55 announcements of the first round of campaigns, only 24 (44%) were carried out in the evenings; the rest were carried out in mornings (20, 36%) or afternoons (11, 20%) in variation with the protocol, and was probably when the vaccinators presumed most people were at home.

##### v. Use of locally made passive cooling devices to store rabies vaccine in wards to support provision of continuous vaccination

To ensure vaccines do not remain outside of the cold chain for more than six months, eight CBC-MDV components were to help to deliver the vaccines to wards in batches. Six out of these were implemented as planned including: coordinated requests and transport systems; basing requests on ward dog population; returning unused vaccines after six months; installation of cooling pots away from sunlight; and monitoring daily temperature in pots. However, labeling of unused vaccines was not carried out as planned; only two out of nine RCs reported having ever returned unused vaccines to the district office and these were given to wards which were not part of the studies for use. Four out of nine pots were not in full use because they developed cracks and leaked when water is added to the cooling sand layer.

The prescribed waste management plans were partially implemented. The different kinds of waste were mostly separated during vaccination clinics (7/9), but instead of returning metallic and biohazard wastes to district offices or nearest health centers for incineration, most teams burnt everything at the location of clinics (6/9), indicating it was safe to do so.

##### vi. A continuous approach to MDV activities; quarterly basis and available on demand by dog owners all year round thereby providing continuous access to dog vaccine

Of the five components of CBC-MDV targeted at supporting provision of continuous dog vaccination, two were implemented as planned, one was modified and two were not implemented as planned. The CBC-MDV protocol prescribed that each strategy team conducts four rounds of campaigns in a year. However, only three out of the nine teams conducted four rounds of campaigns. The frequently cited reasons for variation in vaccination schedules included: farming/ rainy seasons, national activities such as elections, counting of poor households and mass animal vaccination campaigns (in which some RCs participated), social events such as cattle auction days, funerals, puberty rites celebrations and school cycles, with campaigns more patronized on weekends during school terms. For example, some dog owners indicated that during the farming season, either they or their dogs were required in the farms during the day time to guard against monkeys destroying their crops. It was also noted in one district that campaigns were halted during the month-long puberty rites celebrations.

The activity of finding unvaccinated dogs that missed previous rounds was not implemented as planned. The implementers cited that this activity was labor-intensive and not feasible in the absence of an existing village register of dogs. To ensure dog owners have easy and continuous access to vaccinators, the protocol prescribed that OHCs give their mobile numbers out during first round of campaigns. None of OHCs reported giving their numbers out directly as planned (0, 100%) but most (32, 91%) wrote them on the 5-10 posters per village they pasted. The research team observed giving numbers out was practically difficult to do during advertising or vaccination given how busy they were at the centers. However, more than half of OHCs (20, 57%) reported having received calls from dog owners to visit their homes to vaccinate their dogs.

##### vii. Delivery of free dog vaccination clinics using suitable approaches to encourage owner participation

Out of the eight components related to organizing vaccination clinics, five were implemented as planned, one was modified and two were not implemented as planned. The CBC-MDV protocol prescribed that vaccination should take place between 08:00 – 14:00; in practice clinics started as early as 07:00 and as late as 12:00; and closed as early as 11:00 and as late as 18:00. The length of clinics was dependent on turnout at centers. House-to-house campaigns took longer where houses were further apart. The starting time for clinics depended on when farmers had returned home, whether RCs had to perform other duties on the same day (e.g. having to inspect meat) before clinics or whether RCs had to attend to personal business. Vaccinators also cited that microchipping dogs (during which a number of dogs struggled) and entering data into the digital data collection device was time-consuming.

To ensure safe vaccination of dogs by reducing dog aggression, the implementation manual prescribes separation of registration and inoculation points with at least a 20-meter distance and muzzling of potentially aggressive dogs. However, none of the vaccination teams (0/9) implemented these. Dog aggression was associated with poor dog handling techniques by vaccinators. It was observed that vaccinators may not have had enough time to assimilate the benefits of separating dogs being registered and those being inoculated to reduce aggression. Dog aggression was observed to increase the time-per-dog vaccinated and on rare occasions resulted in injury, especially of dog owners.

Muzzles were not used out of fear of being bitten or the muzzles could tear in the process. One rabies coordinator said: “*is too difficult to use muzzles, dogs are too fierce to use it on them, it will get loose, we are afraid, we use the Y-stick”* [RC, Implementation Audit, Strategy 2-Tarime]. Others recommended muzzles of three different sizes, whilst others perceived use of muzzles as time consuming. Consequently, implementers in Butiama and Rorya Districts restrained aggressive dogs by tying the rope or chain on the neck of dogs closely to a tree, and holding the hind legs firmly whilst inoculating the dog. While those in Tarime District used a ‘Y-stick’ to pin down the dog at the neck region with the help of the rope or chain.

The vaccination teams varied the delivery strategies that were prescribed for them, citing the following reasons: villagers saying it was difficult to bring dogs over long distance to centers, large dog populations in their villages, and their own perception of which strategy was likely to reach more dogs. Remarks by implementers indicated they thought subvillage level temporal static point clinics was the most effective approach, with the following quotes exemplifying this, *“subvillage level is very good at reaching more dogs”* [RC, Implementation Audit, Strategy 1-Rorya]; “*the Strategy (subvillage level temporal static point approach) is good because we had time to educate the dog owners”* [RCs, FGD, Strategy 2-Butiama]; *“I think Strategy 3 is good, it covers a lot of places because we use sub-village level (temporal static point approach), house to house and on demand”* [RC, Implementation Audit, Strategy 3-Tarime].

##### viii. Monitoring and feedback on vaccination coverage among stakeholders to promote collaborative local action

Of the six components relating to monitoring, reporting and providing feedback on CBC-MDV, only two (RCs reporting on dogs vaccinated, daily temperature recording of the low-tech cooling devices and rabies events) were delivered as planned. Supervision of campaigns by district veterinary officers was not carried out; the district veterinary officers cited lack of transportation to carry out this task and they expected per diem payment while supervising. OHCs also did not provide weekly reports on dogs needing vaccination and considered the weekly reporting was too frequent to allow for completion. Communities’ self-monitoring of the campaigns and feedback to the research team and the district veterinary office were also not carried out, largely due to weak community involvement in the design of the CBC-MDV and sensitization on this role.

#### Comparing efforts made at delivering CBC-MDV components by strategies Involvement of village level leadership in roll out of CBC-MDV

The strategy teams delivered components relating to involving village leaders with varied degrees of fidelity. For example, very few OHCs discussed their timetables with a village leader to get their approval and support (0/12 for Strategy 1, 2/13 for Strategy 2 and 2/10 for Strategy 3). OHCs largely did not work with ten-cell leaders (Mabalozi) to estimate the dog population in their ward: Strategy 1 (3/12), Strategy 2 (0/13) and Strategy 3 (0/10). Further information about how the delivery of the additional components were delivered is provided in Supplementary Table 2.

All components relating to use of trained village-based OHCs to support vaccination were delivered as planned by all strategies, except sensitization of villagers about campaigns at village meetings which differed: Strategy 1 (2/12), Strategy 2 (8/13) and Strategy 3 (9/10) (suppl table 2). All strategies delivered advertising components as required, but the effort put into the advertising differed: The number of times and hours per village advertised in the first round, and total number of days of vaccination per village were all lowest in Strategy 1 and highest in Strategy 3 respectively (Fig 3). The vaccinators reported that having to walk for a long distance or personally pay for travel by motorbike created challenges to advertising.

**Fig 3.**
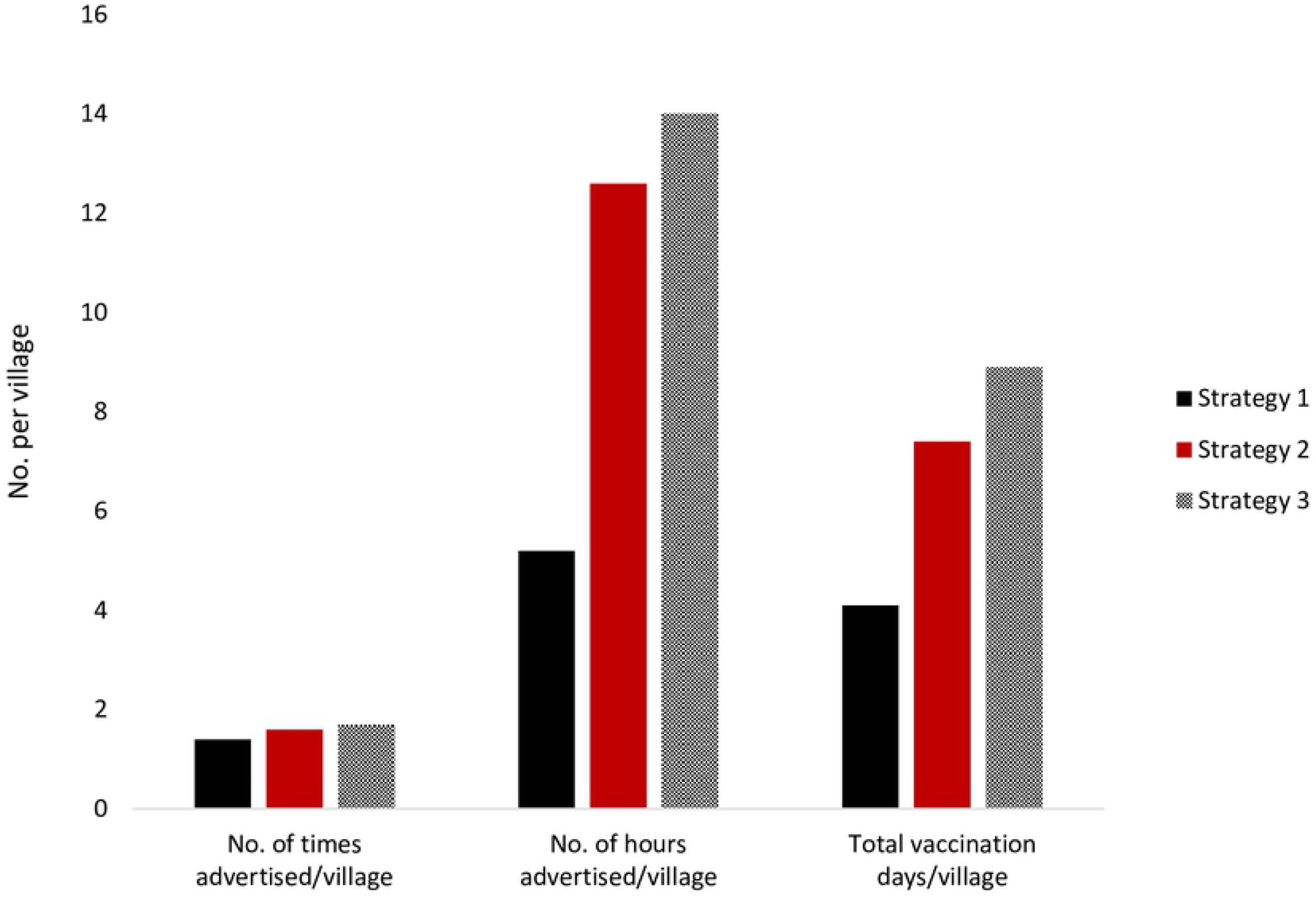
Effort at advertising and delivering vaccination campaigns by strategy (totals for all three team per strategy)

For all strategies, the number of days of campaign activities reduced substantially after the first round. Over the one-year period, the three strategies together used 237 days on campaigns: Strategy 1 (49, 21%), Strategy 2 (95, 40%) and Strategy 3 (91, 39%). The majority of days (189 days, 80%) were spent during the first two rounds (Fig 4).

**Fig 4.**
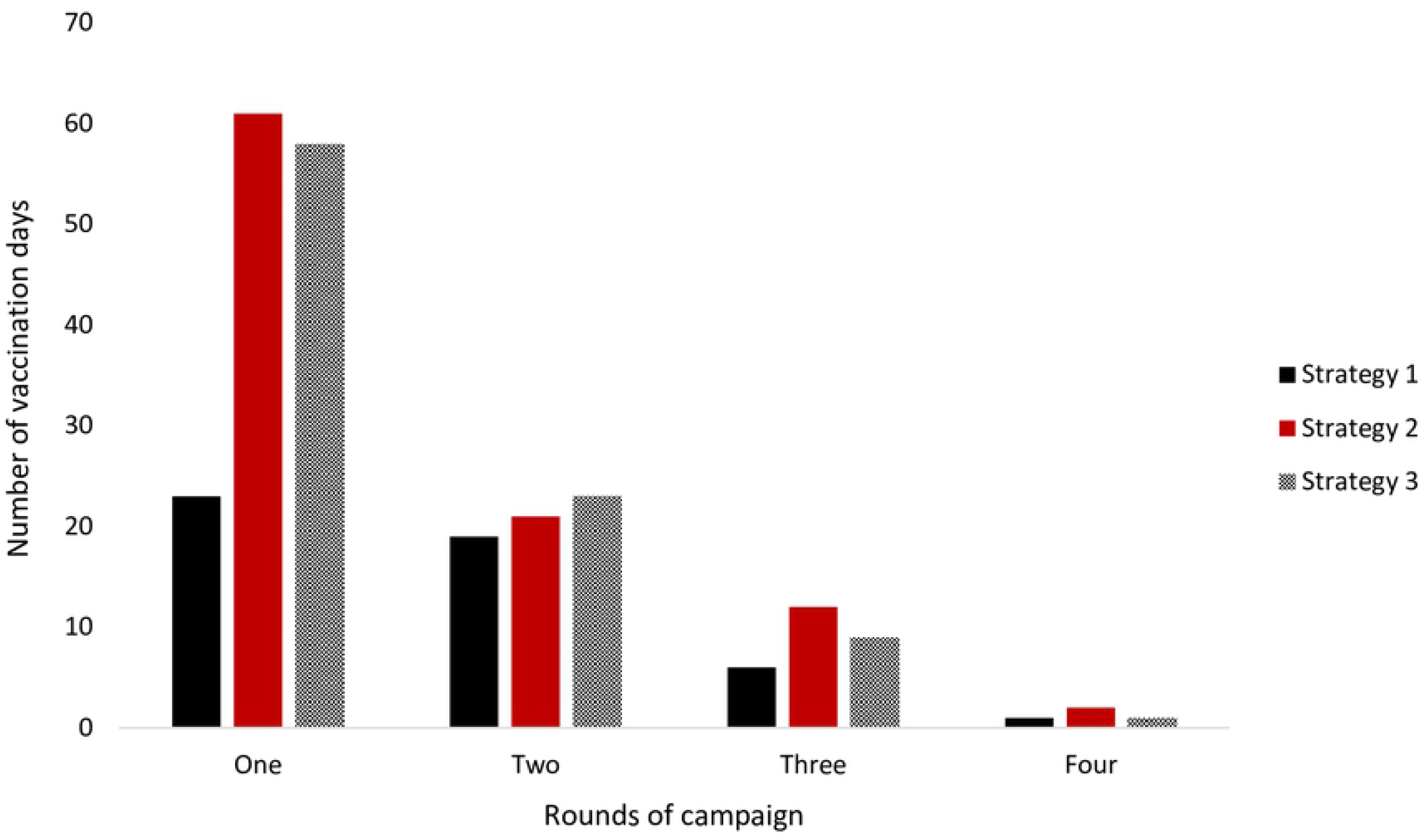
Number of days implemented conducted vaccination activities during each round (totals for all three team per strategy)

The strategy teams differed in terms of numbers of days spent finding dogs that missed central point clinics, responding to on-demand vaccination by dog owners and in organizing quarterly campaigns (suppl table 2).

#### Waste management after vaccination clinic

All teams installed and managed vaccine batches as planned. However, there was discrepancy with regards to how used needles and microchip units were disposed. Some teams either incinerated or disposed of these items in pit toilets: Strategy 1 (2/3 teams), Strategy 2 (2/3 teams) and Strategy 3 (1/3 teams), whilst the rest of the teams burnt all waste at vaccination centers (suppl table 2).

#### Delivery of free dog vaccination clinics using suitable approaches

none of the Strategy teams implemented separating registration and inoculation centers with a distance of at least 20 meters and muzzling of potentially aggressive dogs as planned. The Strategy teams partly followed CBC-MDV manual in selecting approaches to deliver dog vaccination: All Strategy 3 wards opted for subvillage level temporal static point approach, the same approach as was prescribed for use in Strategy 2 wards in round 1 (6/6). In round 2, two of the Strategy 3 wards avoided the lengthy campaign days that come with subvillage level temporal static point approach by deciding to use village level temporal static point. A remark by an RC exemplifies this: “*it (subvillage level temporal static point approach) took long”* [RC, Implementation Audit, Strategy 3-Rorya]. Conversely, two out of the three Strategy 1 teams switched from village level in round 1 to subvillage level temporal static point approach in round 2. The reason given for this switch was that many dogs remained unvaccinated after the round 1 village level temporal static point clinics and so they decided to instead employ a subvillage level temporal static point approach to reach more dogs. All teams employed some house-to-house and on-demand (9/9) approaches. Subvillages were combined for single clinics where implementers considered them to be smaller in size, had smaller dog populations or were closer to each other (suppl Table 3).

Overall, subvillage level temporal static point and on-demand approaches were the most (173 occasions) and least-used (20 occasions) respectively (Fig 5).

**Fig 5.**
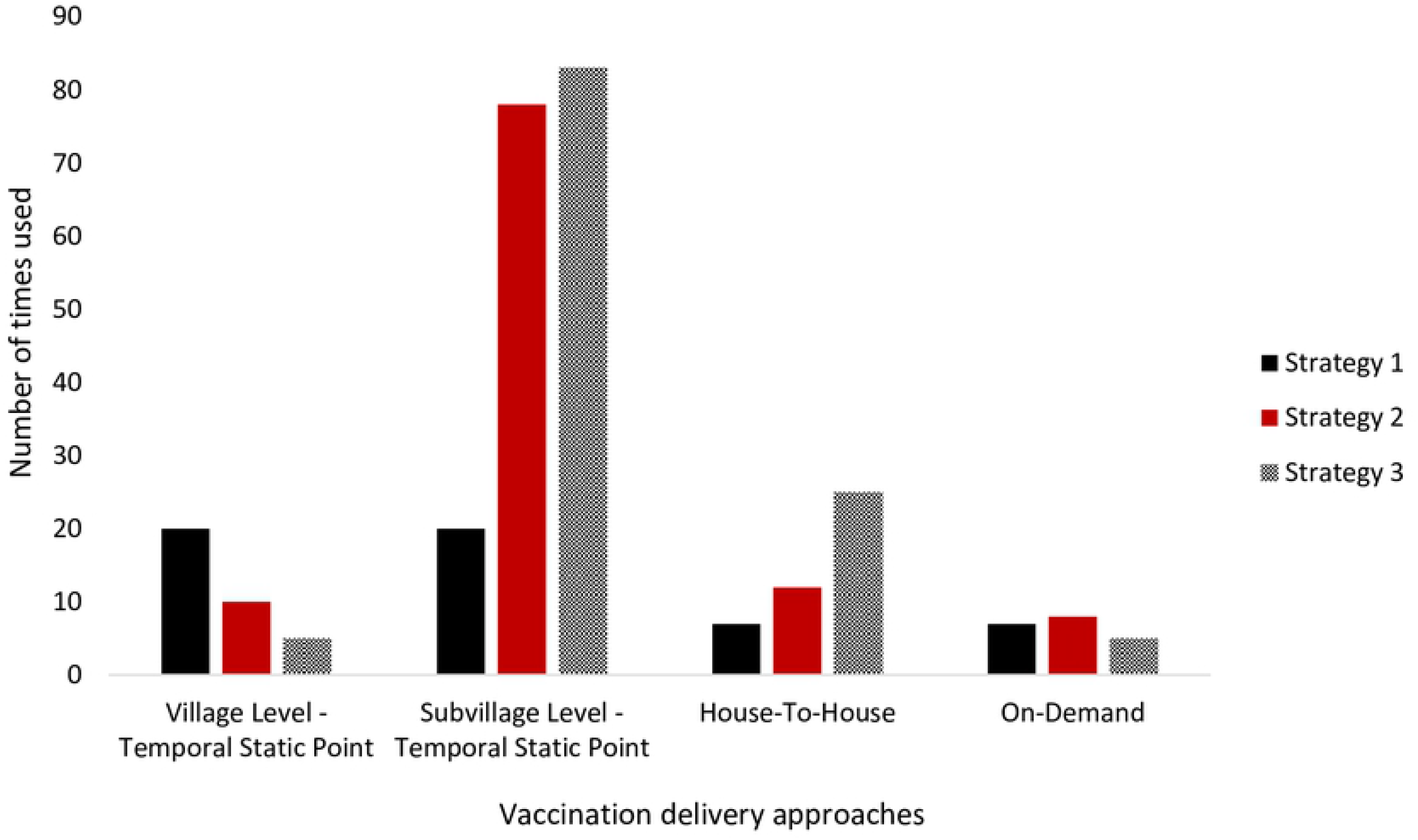
Use of vaccination delivery approaches by strategy team (totals for all three team per strategy)

### Potential effectiveness of the CBC-MDV strategies

To interrupt rabies transmission requires sustaining vaccination coverage above the critical vaccination threshold (approximately 40%). Coverage estimations at month 1 and 11 showed all continuous strategies did sustain coverage above this level, whilst the pulsed approach did not achieve the ≥70% target (Table 5). Coverage at month 11 was slightly lower in Strategy 1 and 3 and slightly increased in Strategy 2, but none were significantly different (Table 5). Strategy 3 which recorded the highest work inputs in terms of advertising and vaccination days, recorded slightly higher annual average vaccination coverage: Strategy 1, 2 & 3 (61.43%, 62.93% & 63.46%), respectively (Table 5).

**Table 5.** Vaccination coverage achieved by the delivery strategies at month 1 and 11

| Vaccination coverage achieved by delivery strategies |  |  |  |
| --- | --- | --- | --- |
| Strategies Arms | Month – 1 (%) | Month – 11 (%) | Annual Averages (%) |
| Pulse | 35.86 | 32.10 | 33.98 |
| Strategy 1 | 65.07 | 57.78 | 61.43 |
| Strategy 2 | 60.97 | 64.88 | 62.93 |
| Strategy 3 | 68.00 | 58.91 | 63.46 |

### Optimization of CBC-MDV for replication in the full-scale trial and dissemination in other contexts

Table 6 details optimization of some components of CBC-MDV for replication in the full-scale trial and lessons for dissemination in other contexts.

**Table 6.**
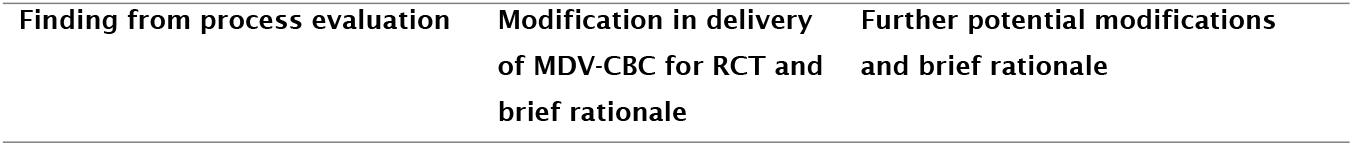
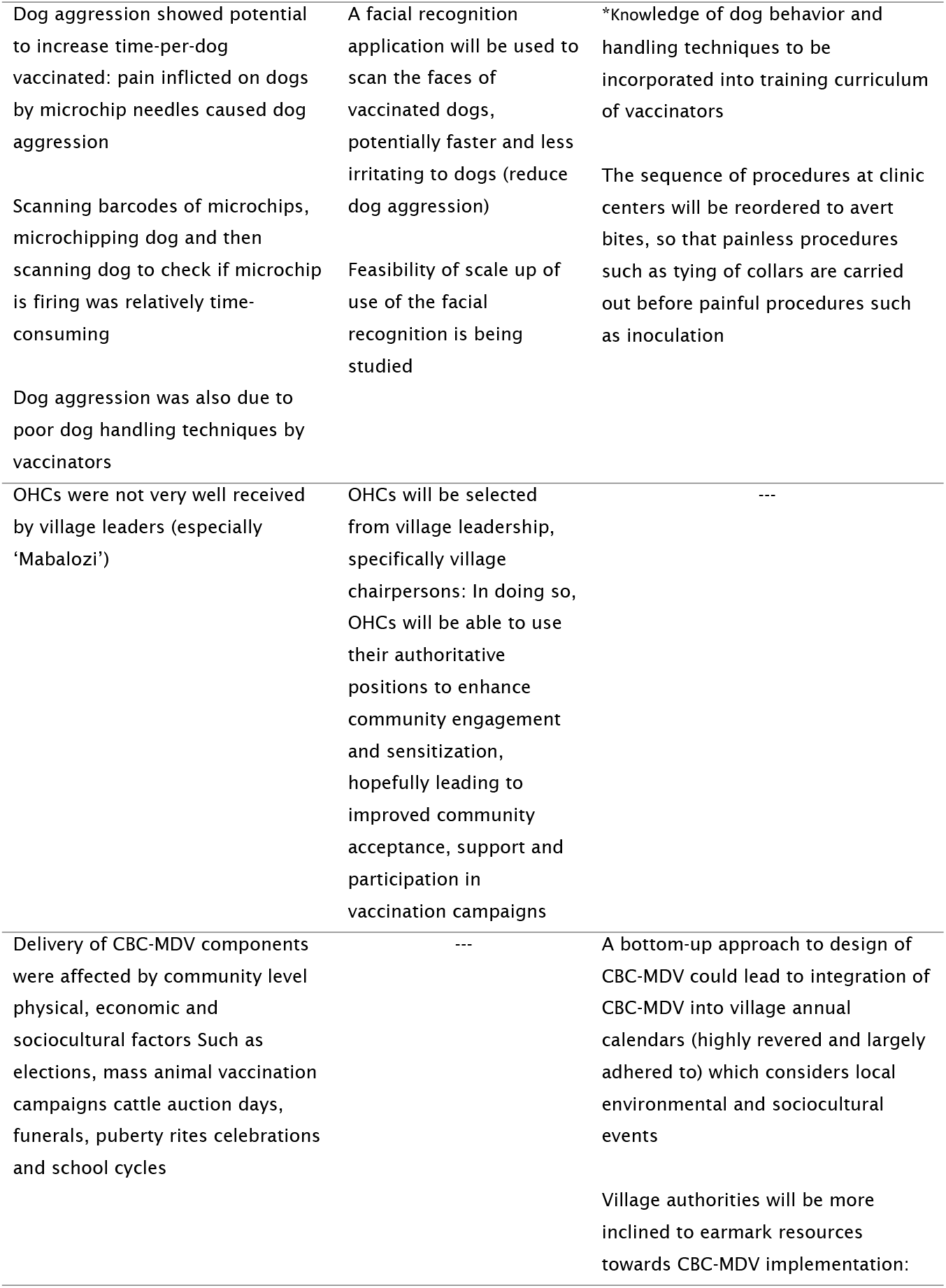
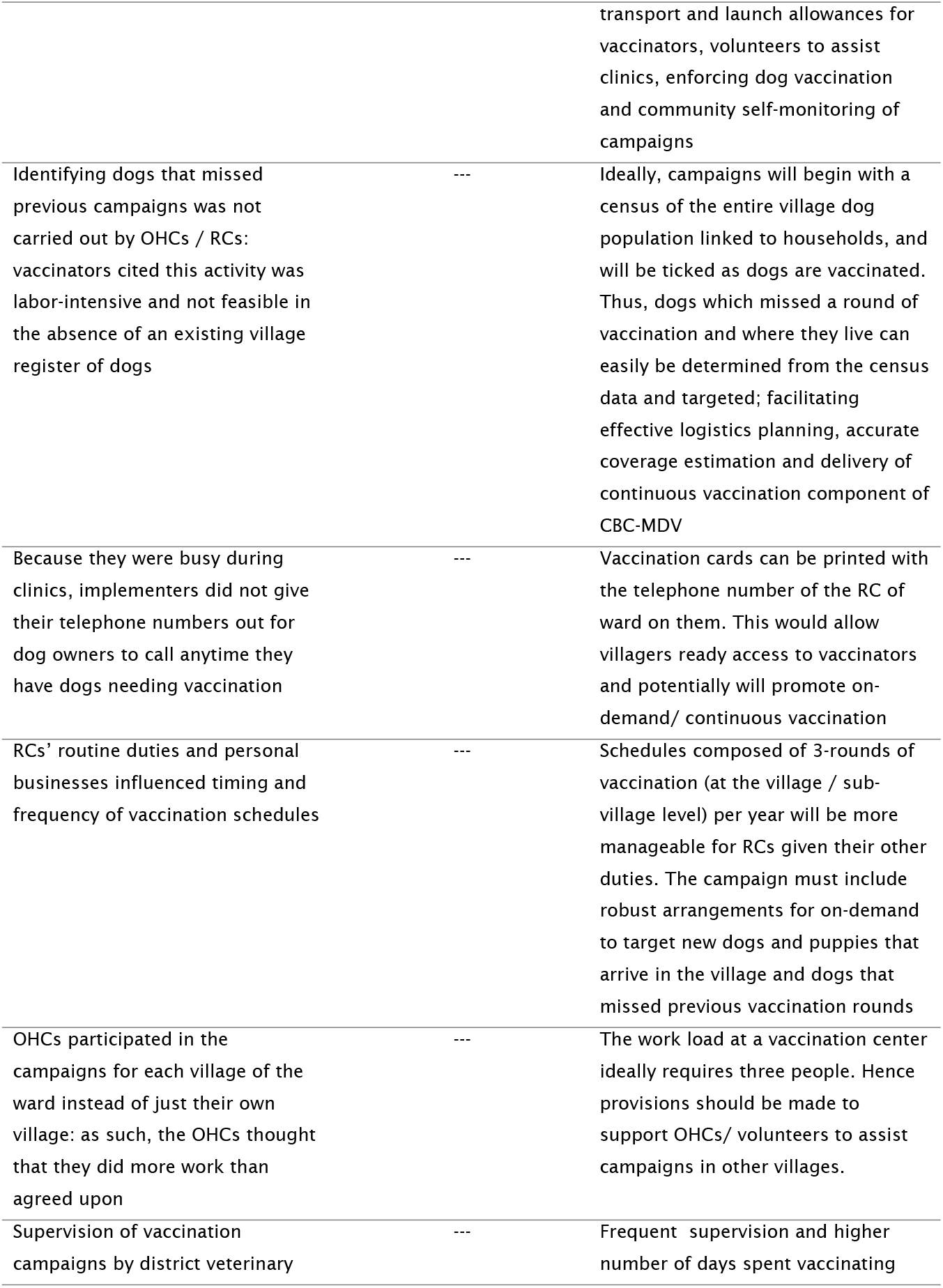
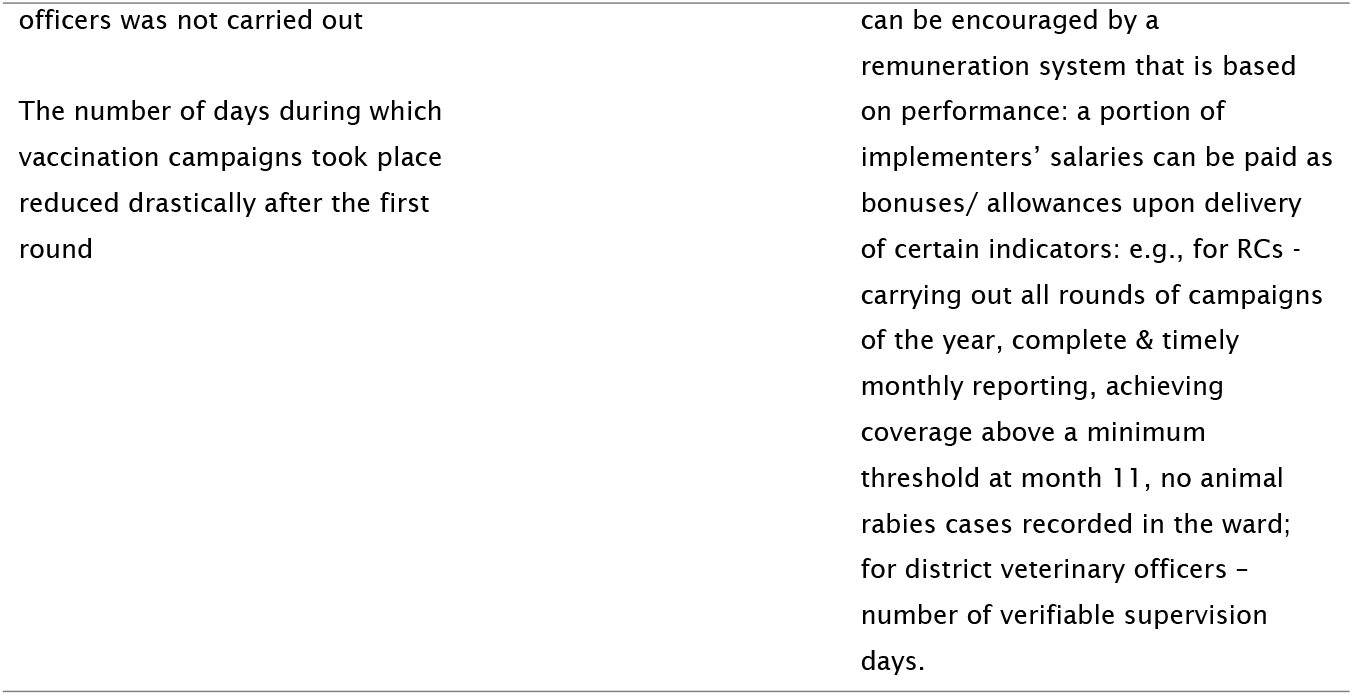
How CBC-MDV can be optimized for replication in the full-scale trial and dissemination in other contexts

## Discussion

This paper provides formative insights into the development, feasibility, potential effectiveness and optimization of CBC-MDV. Key findings were: *i*. the development process of CBC-MDV was iterative and involved stakeholders from multiple sectors but did not include direct involvement of community members; *ii*. It was feasible to deliver about half of CBC-MDV components as planned (about 50% fidelity to implementation manual); *iii*. variation of delivery from what the implementation manual prescribed was because of factors inherent in the design of the CBC-MDV strategies, implementers’ understanding and appreciation of the CBC-MDV components and moderating effects of contextual (sociocultural, economic, political and environmental) elements such as elections, mass cattle vaccination campaigns, cattle auction days, funerals, puberty rites celebrations and school cycles; *iv*. all continuous delivery strategies of CBC-MDV sustained vaccination coverage above the critical threshold (approximately 40%), whilst the pulse (once annual) strategy failed to achieve the required ≥70% vaccination coverage; and *v*. because of the variation to protocols a number of CBC-MDV components needed optimization prior to replication in the planned full-scale trial.

The absence of community involvement in the design stage of CBC-MDV and weak community sensitization at roll out may explain why some village leaders perceived the project as an avenue for making money, questioned the identity of OHCs or didn’t fully corporate. More effective community entry processes could have enhanced participation, strengthened collaborations between implementers and community leaders in mobilizing towards vaccination campaigns and increase community support and contributions to implementation [24–28]. Globally, community participation in intervention delivery has evolved from communities as passive recipients, through communities as active participants in delivery to communities as co-designers of interventions [16,25]. The performance of the community-based personnel in the delivery of CBC-MDV components and outcomes of community-led deliveries of interventions elsewhere show that communities can implement intervention such as dog vaccination campaigns with effective engagement and if supplied with logistics [25,29].

Implementation of CBC-MDV components that relate to managing vaccination logistics, organizing clinics and information recording were carried out with high fidelity. On the other hand, components aimed at ensuring vaccination clinics proceeded smoothly such as community engagement and sensitization, supervision and monitoring of campaigns, separation of registration and inoculation points to minimize dog aggression were mostly omitted or implemented with low fidelity. Certain components such as finding dogs that missed previous rounds, vaccinators giving their telephone numbers out to dog owners at centers and muzzling of potentially aggressive dogs appeared practically challenging to implement. For instance, some implementers expressed fear about muzzling a dog, others indicated the muzzles were too small or could tear in the process. As documented by other process evaluation studies, implementers not having ample time to assimilate the value of each intervention component, to schedule activities, not feeling competent enough to deliver certain components or having unusable equipment resulted in low fidelity [30,31].

The marked decline in the number of vaccination days with each round of vaccination may be an indication of implementation fatigue. RCs serve large populations (3-4 villages / ward on average) providing many different extension services such as dipping of large herds of domestic animals, meat inspection at several different locations, animal levy collection at cattle auctions and other routine duties. It is likely that conducting four rounds of dog vaccination campaigns was a substantial additional work burden. It is also possible that the RCs did not consider the *continuous* component of CBC-MDV as critical, and rather assumed that they had vaccinated sufficient dogs in Round 1 without consideration of arrival of new dogs and puppies in villages. This would be consistent with other studies that found staff ‘burn out’ as a barrier to implementing community-based interventions as intended [32,33].

Variation in work inputs by Strategy teams were explained primarily by the nature of the specific approaches and offer insights into variation in the coverage by each strategy. Strategy 1 required a larger effort occurring over a shorter period of time for the implementers. However, because the vaccination activity of Strategy 1 occurs at a central point of the village, for many owners this Strategy likely posed a challenge of access as they will be required to travel further to reach the central point where the clinic was hosted. Living far from the point of the clinic is commonly cited as a reason for nonparticipation in other studies [34–37]. In comparison, Strategy 2, being hosted at the subvillage level, comes with a relatively lighter workload on each vaccination day for the RC, but, with multiple subvillages for every village, requiring multiple days to complete the campaign (reaching 35 consecutive days). However, subvillage level clinics are easier for the owners to attend. It is noteworthy that, when given the discretion to choose, all Strategy 3 teams adopted the subvillage (Strategy 2) approach even though they reported it requires more time. This suggests that empowering implementers with the ability to choose their own Strategy fostered a stronger sense of ownership and desire to do what it takes to achieve more. This notion is supported in previous research where social motivation has been reported to enhance community participation in community level development activities [38]. The discretion also may have allowed Strategy 3 teams to be more flexible in their schedules around personal and local events.

Strategy 3 teams also recorded higher number of times and hours advertised per village and number of vaccination days per village, and possibly explain why the annual average vaccination coverage achieved by Strategy 3 was marginally higher (Lugelo et at., preprint). However, the discretion may have caused strategy 3 team to relax after the first round of clinics, hence they account for 3 out of the 7 missed rounds by all strategies and recording a lower coverage at month 11. Given the differences in the prescribed activities, it seems logical that Strategy 1 teams would need to work harder during subsequent rounds to attain similar outputs as strategies 2 and 3.

Including communities in evaluating outcomes of CBC-MDV is likely to foster ownership and sustained efforts at delivering components. Community participation in evaluating local interventions has been gaining traction and, for example, was a key component of the community-directed treatment with ivermectin (CDTI) model introduced by the African Programme for Onchocerciasis Control [16,25]. In the CDTI model, a 3-member committee selected by each village carried out community self-monitoring of mass distribution of ivermectin, thereby checking the performance of distributors and compliance of community members. In the process, challenges were identified and resolved with participation of community leaders. Lessons and strategies such as those outlined above and those generated from this study could be incorporated into CBC-MDV to ensure its successful replication.

Several local environmental, economic and sociocultural events affected feasibility of delivering CBC-MDV components. Structural community participation in initializing and implementing the intervention would clearly be valuable in identifying these events and issues, and replication of CBC-MDV across wider contexts would benefit from tailoring campaign schedules to local environmental and social events [36,37,39,40] or calendars.

Process evaluation has been carried out for a wide range of complex interventions, but this study represents the first process evaluation of mass dog vaccination campaigns to our knowledge. The study revealed implementation bottlenecks in their delivery, understanding of the impact pathways underpinning these bottlenecks and also opportunities for addressing them. These insights could be of value when designing national rabies elimination strategies.

The study is likely slightly affected by recall bias where data collection processes depended to a large extent on implementer reports. However, use of mixed methods, including non-participant observations and following the intervention prospectively through the design and implementation phases provided first hand observations.

## Conclusions

The development of CBC-MDV incorporated extensive stakeholder views and hence fostered strong stakeholder acceptance so far. Including community level decision makers/ leaders would be likely to have fostered ownership among communities as well. It is possible to delivery CBC-MDV and achieve good vaccination coverage although intervention-, implementer- and context-related factors influenced delivery CBC-MDV components and effectiveness of the strategies at reaching more dogs. These factors altogether occasioned variations in implementation of about half of CBC-MDV components, resulting in differences in choices and outputs made by strategy teams. The CBC-MDV strategies sustained vaccination coverage well above the critical threshold (approximately 40%) throughout the year whilst the pulse Strategy failed to achieve the required vaccination coverage of ≥70%. The findings are being used to optimize CBC-MDV components for dissemination in an RCT across the entire Mara region. Overall, we conclude that improved supervision and monitoring, as well as community participation in design, planning and execution of MDV could result in higher fidelity, dose and reach of CBC-MDV strategies in a more sustainable manner.

## Author contributions

**Conceptualization:** Christian Tetteh Duamor, Katie Hampson, Felix Lankester, Sally Wyke, Sarah Cleaveland.

**Data collection:** Christian Tetteh Duamor, Ahmed Lugelo.

**Formal analysis:** Christian Tetteh Duamor, Ahmed Lugelo.

**Funding acquisition:** Katie Hampson, Felix Lankester, Sally Wyke, Sarah Cleaveland.

**Investigation:** Christian Tetteh Duamor.

**Methodology:** Christian Tetteh Duamor, Sally Wyke.

**Project administration:** Christian Tetteh Duamor, Ahmed Lugelo, Felix Lankester.

**Supervision:** Katie Hampson, Felix Lankester, Katharina Kreppel, Emmanuel Mpolya, Sally Wyke, Sarah Cleaveland.

**Validation:** Christian Tetteh Duamor, Katie Hampson, Felix Lankester, Katharina Kreppel, Sally Wyke, Sarah Cleaveland.

**Visualization:** Christian Tetteh Duamor.

**Writing – original draft:** Christian Tetteh Duamor.

**Writing – review & editing:** Christian Tetteh Duamor, Katie Hampson, Felix Lankester, Katharina Kreppel, Emmanuel Mpolya, Sally Wyke, Sarah Cleaveland.

---

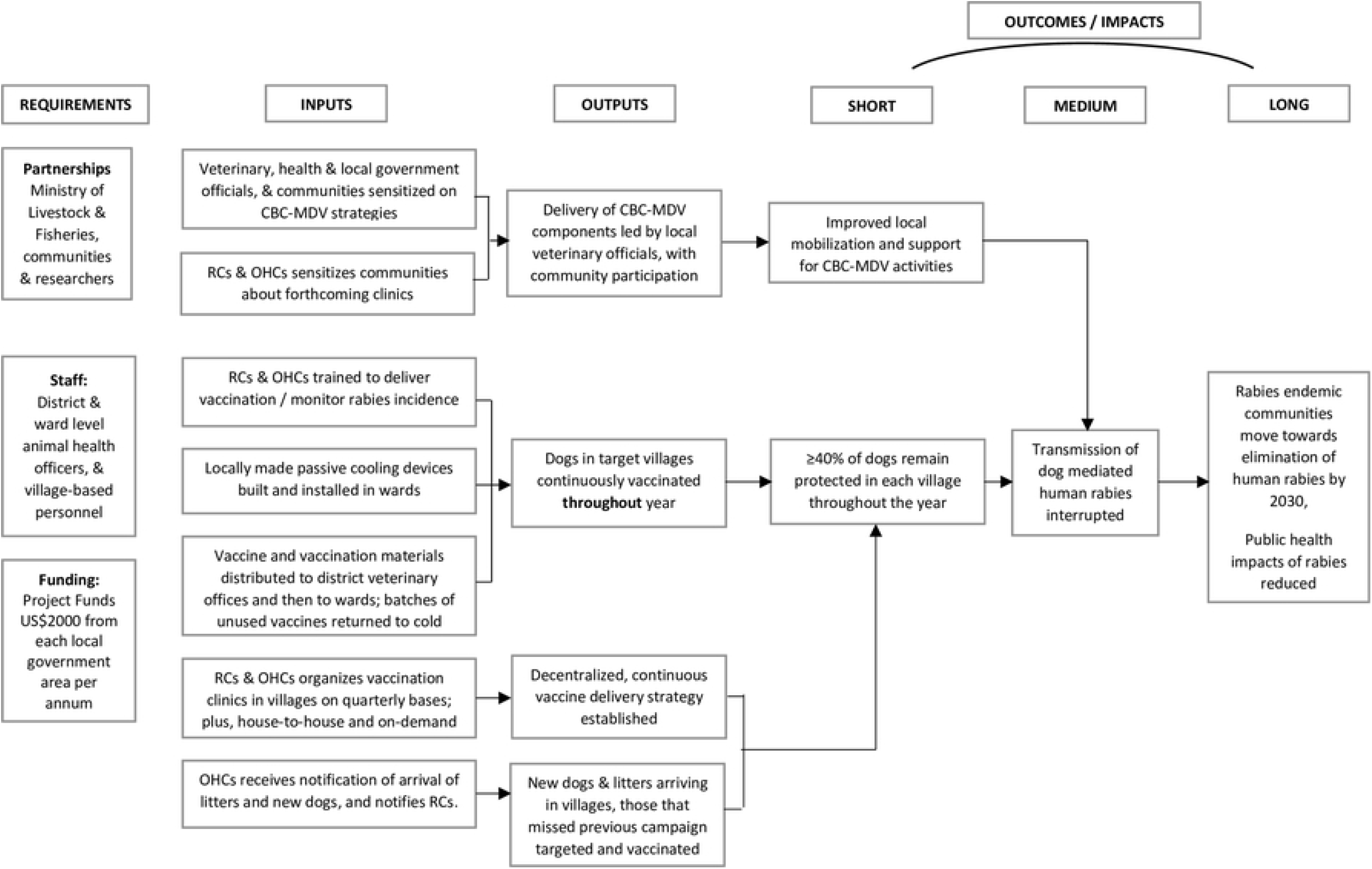

